# Rejuvenation-Responsive and Senolytic-Sensitive Muscle Stem Cells Unveiled by CD200 and CD63 in Geriatric Muscle

**DOI:** 10.1101/2025.08.29.673011

**Authors:** Ye Lynne Kim, Young-Woo Jo, Takwon Yoo, Kyusang Yoo, Ji-Hoon Kim, Myungsun Park, In-Wook Song, Hyun Kim, Yea-Eun Kim, Sang-Hyeon Hann, Jong-Eun Park, Daehyun Baek, Young-Yun Kong

## Abstract

Muscle stem cells (MuSCs) are parenchymal cells in skeletal muscle regeneration and maintenance. With aging, MuSCs experience a decline in their regenerative function and reduction in their number. However, recent evidence points to substantial heterogeneity within the aged MuSC population, raising questions about the underlying mechanisms of age-associated dysfunction. Here, we used Pax7^CreERT2^;Rosa^YFP^ mice (MuSC^YFP^) to label Pax7-expressing MuSCs and chronologically traced MuSCs until geriatric age. Genetic labeling and chronological tracing revealed that the number of YFP^+^ MuSC remained comparable between young, middle and geriatric ages. At geriatric age, YFP^+^ MuSCs exhibited reduced expression of traditional MuSC markers such as VCAM1 and PAX7. A previously unrecognized subpopulation emerged, characterized by loss of VCAM1 and low or absent PAX7. Despite their altered marker profile, these cells retained transcriptional signatures of quiescence and myogenic potential, but displayed significantly reduced proliferative and regenerative capacities. They displayed gene expression patterns indicative of senescence-like state and were selectively ablated by senolytic treatment. DHT restored regenerative function in aged mice and re-induced VCAM1 expression in YFP^+^/Pax7^-/low^/VCAM1^-^ cells, indicating responsiveness to rejuvenation. Based on their emergence with aging, functional impairment and responsiveness to rejuvenation, we termed this population GERI-MuSCs (Geriatric Emerging Rejuvenation-responsive and Impaired MuSCs). CD63 and CD200 were identified as novel surface markers that together with VCAM1, reliably detect GERI-MuSCs as well as classical Pax7^+^/VCAM1^High^ MuSCs, providing a tool for comprehensive isolation of MuSCs from aged wild-type mice. Together, our findings provide a refined framework for studying MuSC aging and offer new tools for isolating functionally distinct MuSC subsets from aged skeletal muscle.

## Introduction

Skeletal muscle regeneration is orchestrated primarily by tissue resident stem cells, known as muscle stem cells (MuSCs) or satellite cells(Mauro, 1961). MuSCs reside beneath the basal lamina of myofibers in a quiescent state under homeostatic condition but become rapidly activated upon injury to proliferate, differentiate, and fuse into regenerating myofibers(Evano and Tajbakhsh, 2018). While the regenerative function of MuSCs is well maintained in youth, a decline in their number and regenerative competence is observed in aging, leading to impaired muscle repair and tissue maintenance(Blau et al., 2015).

The age-related decline in MuSC function arises from a combination of extrinsic and intrinsic factors. Extrinsic factors include the increased pro-inflammatory cytokines(Degens, 2007), extracellular matrix remodeling(Schuler et al., 2021) and altered mechanical properties of the niche microenvironment(Chakkalakal et al., 2012). MuSC intrinsic factors include disrupted metabolic homeostasis(Pala et al., 2018), loss of proteostasis(Kim et al., 2021b), altered epigenetic regulation(Hernando-Herraez et al., 2019), mitochondrial dysfunction(Rygiel et al., 2016) and aberrant activation of signaling pathways, including p38 MAPK(Bernet et al., 2014), JAK/STAT(Price et al., 2014) and mTORC1(Tang et al., 2019). As a result, aged MuSCs show reduced colony-forming capacity, failure to maintain quiescence and loss of self-renewal potential causing a significant decline in the MuSC pool. This reduction in the resident stem cell population directly undermines the muscle’s capacity for effective repair(Blau et al., 2015, Day et al., 2010). Although functional and molecular differences between young and aged MuSCs have been extensively documented, the timing and trajectory by which these alterations arise remain unclear. This study was therefore designed to delineate the transition that occurs between middle-aged and geriatric MuSCs, rather than to re-characterize differences that are already well established between young and aged muscles.

Efforts to reverse MuSC dysfunction have explored both systemic and cell-intrinsic interventions. Circulating factors identified in parabiosis experiments(Conboy et al., 2005), as well as supplementation with molecules such as NAD^+^(Zhang et al., 2016) and oxytocin(Elabd et al., 2014), have shown potential in restoring regenerative capacity. Pharmacological modulation of signaling pathways, such as inhibition of JAK/STAT(Price et al., 2014), p38 MAPK(Bernet et al., 2014), mTORC1, TGF-β or Wnt signaling(Carlson et al., 2009), has led to improved regeneration in aged muscle. However, variation in the age of experimental animals complicates the interpretation of these findings. Many rejuvenation studies define ‘aged’ mice as 18 to 24 months old. This range encompasses a period of heterogeneous functional decline, during which a substantial fraction of MuSCs may still retain regenerative capacity. Therefore, since it is unclear whether these models fully represent a truly geriatric state, questions remain as to whether these interventions directly restore dysfunctional MuSCs, selectively expand a subset of aged MuSCs that retain partial functionality, or delay or prevent the aging of MuSCs that have not yet undergone substantial aging. To address this uncertainty, we chronologically traced the fate of Pax7-expressing MuSCs in geriatric mice using a genetic labeling system. This allowed us to evaluate the composition, functionality, and responsiveness of aged MuSC subsets.

In addition to interventions aimed at reversing MuSC dysfunction, an alternative strategy to restore regenerative competence involves targeting senescent cells. Senescent cells accumulate in aged muscle and are thought to impair regeneration through secretion of pro-inflammatory and matrix-remodeling factors, collectively known as the senescence-associated secretory phenotype (SASP) (Saito and Chikenji, 2021). Senolytic agents, such as dasatinib and quercetin, which selectively eliminate senescent cells have been shown to improve tissue homeostasis and extend muscle healthspan in preclinical models(Xu et al., 2018). However, although senescence is broadly defined as a state of irreversible cell cycle arrest accompanied by transcriptional reprogramming, how this process manifests in quiescent adult stem cells *in vivo* remains incompletely understood. This uncertainty raises concerns regarding the use of senolytics, as such treatment may inadvertently eliminate MuSC subsets that retain latent regenerative potential. A better understanding of how senescence-like features emerge within the aged MuSC pool is therefore critical for evaluating the risks and benefits of senolytic treatment in aging muscle.

In this study, our findings reveal an age-dependent accumulation of a previously unrecognized population of MuSCs, which we termed GERI-MuSCs (Geriatric Emerging Regeneration-capable and Impaired MuSCs). GERI-MuSCs are characterized by low expression of traditional marker genes, impaired yet retained functionality, susceptibility to senolytic deletion, and responsiveness to DHT rejuvenation. Moreover, CD63 and CD200 can be reliable markers that can be used with VCAM1 to detect previously hard-to-detect MuSCs. This combinatorial strategy provides a framework for future isolation and therapeutic targeting of GERI-MuSCs. Together, our results refine the understanding of in vivo MuSC aging and highlight the importance of studying stem cell subsets that are functionally impaired yet not irreversibly senescent.

## Results

### Genetic labeling of Pax7^+^ cells reveal marker-deficient MuSCs in aged muscle

*Paired box protein 7* (*Pax7*) is a transcription factor widely recognized as a defining marker of MuSCs (Seale et al., 2000). To investigate how aging alters the identity of MuSCs, we used *Pax7^CreERT2^;Rosa^YFP^*mice (MuSC^YFP^). In this model, tamoxifen administration at 3-5 months of age irreversibly labels Pax7-expressing MuSCs. YFP^+^ cells were isolated at young (3-5 months), middle (10-14 months) and geriatric (>28 months) ages, using a conventional surface markers and gating strategy: stem cell antigen-1 negative (SCA-1^−^) and vascular cell adhesion protein 1 positive (VCAM1^+^) cells among lineage negative (CD31^−^/CD45^−^) and live cells (Liu et al., 2015) (Fig. 1A). As previously reported, the number of VCAM1^+^/SCA-1^−^ MuSCs declined significantly with age, from an average of 12.22% at young and 13.62% at middle age to 8.31% at geriatric age (Fig. 1B). The variability of VCAM1^+^ MuSC number at geriatric age likely reflects mouse-to-mouse differences in the onset and progression of late-life phenotypes. Despite this heterogeneity, the overall shift toward a VCAM1-low state was observed across all geriatric replicates. Interestingly, however, when MuSCs were defined by YFP expression alone, their number remained comparable across ages (Fig. 1C). The broader spread observed in the young group likely reflects the known heterogeneity of MuSCs in early adulthood, which encompass multiple transcriptional and functional substates (Dong Seong Cho, 2017). This developmental heterogeneity results in greater variability in young mice compared to the more uniform profiles seen in middle-aged. Although the young group displayed greater biological variability, the overall mean MuSC frequencies were comparable among age groups.

**Figure. 1.**
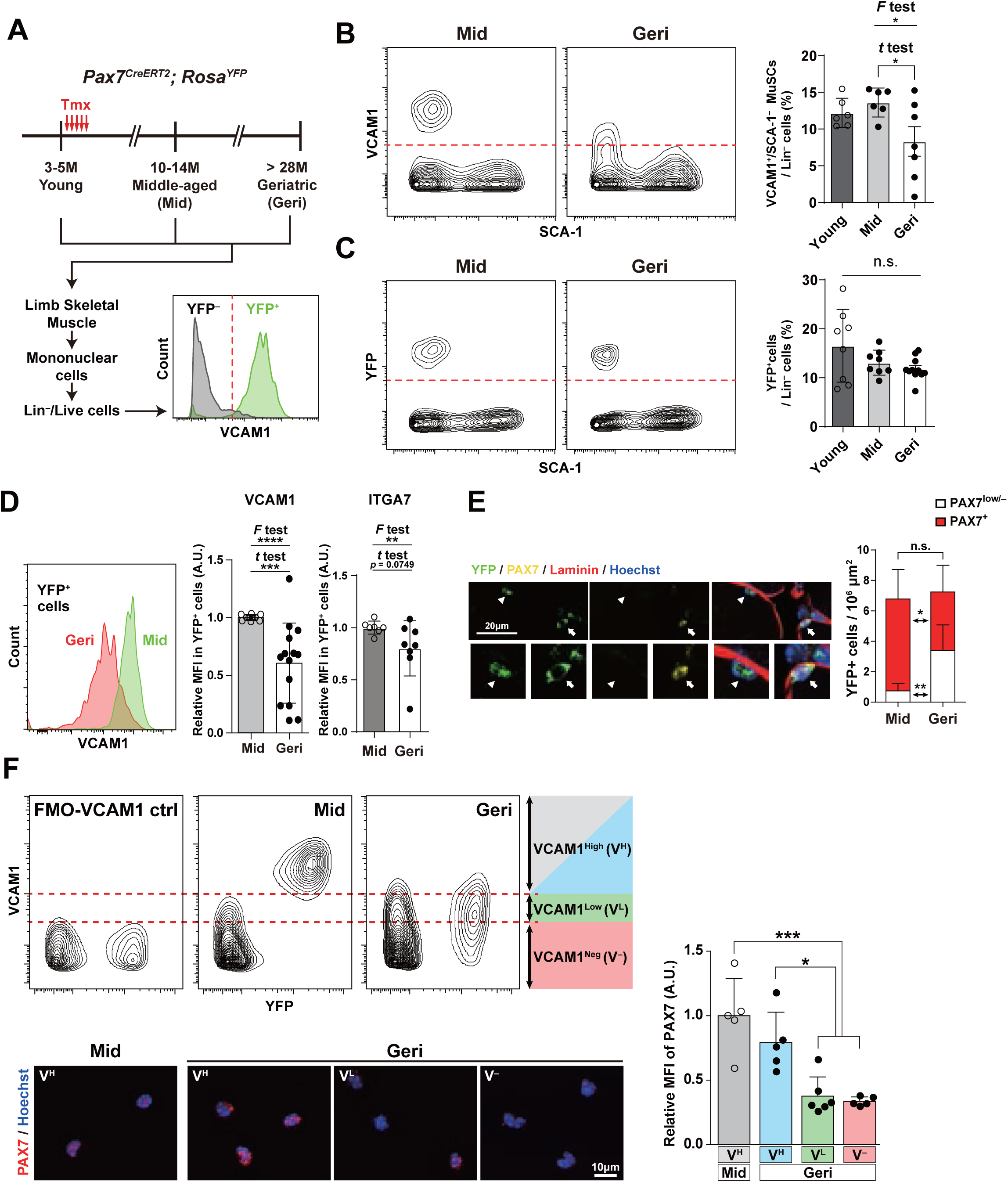
The YFP^+^ cells from *Pax7^CreERT2^;Rosa^YFP^* mice do not decrease with age. (**A**) Scheme of MuSC labeling in *Pax7^CreERT2^;Rosa^YFP^* mice (MuSC^YFP^) and analysis at middle (Mid) and geriatric age (Geri) using fluorescence-activated cell sorting (FACS) strategy for isolation of YFP^+^ MuSCs using traditional marker, VCAM1. (**B, C**) Representative plot and quantification for VCAM1^+^ MuSCs (**B**) and YFP^+^ cells (**C**) in lineage negative (Lin^−^) cells of Young (n=6 and n=8 each), Mid (n = 7, n = 8, each) and Geri (n = 6, n = 11, each) MuSC^YFP^, Welch’s *t* test and *F*-test; *p* = 0.0441 and 0.0460 for (**B**), respectively. (**D**) Representative FACS histogram and quantification of VCAM1 and ITGA7 expression in YFP^+^ cells of Mid and Geri MuSC^YFP^. Quantification of relative VCAM1 and ITGA7 mean fluorescence intensity (MFI) in YFP^+^ cells of Mid (n = 12, n = 6, each) and Geri (n = 14, n = 8, each) MuSC^YFP^, Welch’s *t* test and *F*-test; *p* = 0.0009 and <0.0001. (**E**) Representative images and quantification of immunohistochemistry staining for Hoechst (Blue), YFP (Green), PAX7 (Yellow), and Laminin (Red) in the *Tibialis Anterior* (TA) of MuSC^YFP^. Arrows indicate PAX7^+^/YFP^+^ cells and arrowheads indicate PAX7^−^/YFP^+^ cells under basal lamina. Quantification of the number of Pax7^+^/YFP^+^ and PAX7^−^/YFP^+^ cells from IHC staining of TA in Mid (n = 7) and Geri (n = 9) MuSC^YFP^, two-way ANOVA followed by Bonferroni’s multiple comparison test, PAX7^+^/YFP^+^; *p* = 0.0205, PAX7^-^/YFP^+^; *p* = 0.0051. (**F**) Division of YFP^+^ cells according to their VCAM1 MFI as VCAM1^High^ (V^H^), VCAM1^Low^ (V^L^), and VCAM1^Neg^ (V^−^) followed by representative images and quantification of immunocytochemistry (ICC) staining for Hoechst (Blue), PAX7 (Red) of YFP^+^ cells. VCAM1 low and negative MuSCs represent approximately 44.6% ± 35.7%, whereas VCAM1 high MuSCs represent 3.9% ± 1.8%. Quantification of PAX7 fluorescence intensity from isolated YFP^+^ cells of Mid V^H^, Geri V^H^, V^−^ (n = 5, each), and V^L^ (n=6), one-way ANOVA followed by Bonferroni’s multiple comparison test; *p* = 0.0004 (vs Mid V^H^), and 0.0179, 0.0123 (Geri V^H^ vs V^L^, V^−^, respectively).

To investigate the disparity in the numbers of surface marker labelled MuSCs and genetically labelled MuSCs at geriatric age, we examined the expression of traditional MuSC surface markers, VCAM1 and integrin alpha 7 (ITGA7) in YFP^+^ population (Blanco-Bose et al., 2001, Sacco et al., 2008). We observed a significant decline in VCAM1 expression, while ITGA7 expression remained comparable (Fig. 1D). Notably, both VCAM1 and ITGA7 expression showed significantly greater variation at geriatric age, in YFP^+^ cells, likely due to inter-individual variability during aging. In young and middle-aged muscle, the VCAM1^low^ and VCAM1^-^ MuSC populations were extremely small and often nearly absent, indicating that these subsets may be not a substantial component of the MuSC pool earlier in life. The detectable emergence and expansion of VCAM1^low^ cells occurred predominantly at the geriatric stage, suggesting that this population represents an aging-associated state rather than a pre-existing subset.

Since YFP^+^ cells at geriatric age have varying marker gene expression, we investigated their localization *in vivo* by conducting immunohistochemistry of *tibialis anterior* (TA) muscle sections for PAX7. YFP^+^ cells showed sublaminar localization and co-expressed PAX7 in middle aged mice (Fig. 1E). In geriatric aged mice, however, the number of PAX7^+^/YFP^+^ cells decreased, while YFP^+^/PAX7^low/–^ cells concomitantly increased (Fig. 1E).

To examine the relationship between PAX7 and varying MuSC marker expression, YFP^+^ cells were grouped by VCAM1 expression levels: VCAM1^High^, VCAM1^Low^, and VCAM1^−^, and immunostained for PAX7. The PAX7 fluorescence intensity was comparable between VCAM1^High^ cells from middle-aged and geriatric mice, but was markedly reduced in VCAM1^Low^ and VCAM1^−^ cells of geriatric mice (Fig. 1F). Together, these findings show an emerging population of YFP^+^ cells that lose traditional markers like Pax7 and VCAM1 at geriatric age.

### YFP^+^/VCAM1^−^ cells are quiescent MuSCs

To further explore the characteristics of YFP^+^/VCAM1^−^ cells in geriatric mice, we examined their potential myogenic status. Given that loss of canonical markers might reflect lineage progression, we assessed the expression of myogenin (MYOG), a marker of late-stage myogenic differentiation, in YFP^+^ cells from uninjured middle, geriatric ages and 5-days post injury. We found that MYOG expression was largely absent in YFP^+^ cells from uninjured aged muscle, even in geriatric YFP^+^/VCAM1^Low^ and YFP^+^/VCAM1^−^ cells (Fig. 2A-C). This contrasts with robust MYOG signal detected at 5 days post injury, suggesting that YFP^+^ cells are not committed myoblasts.

**Figure 2.**
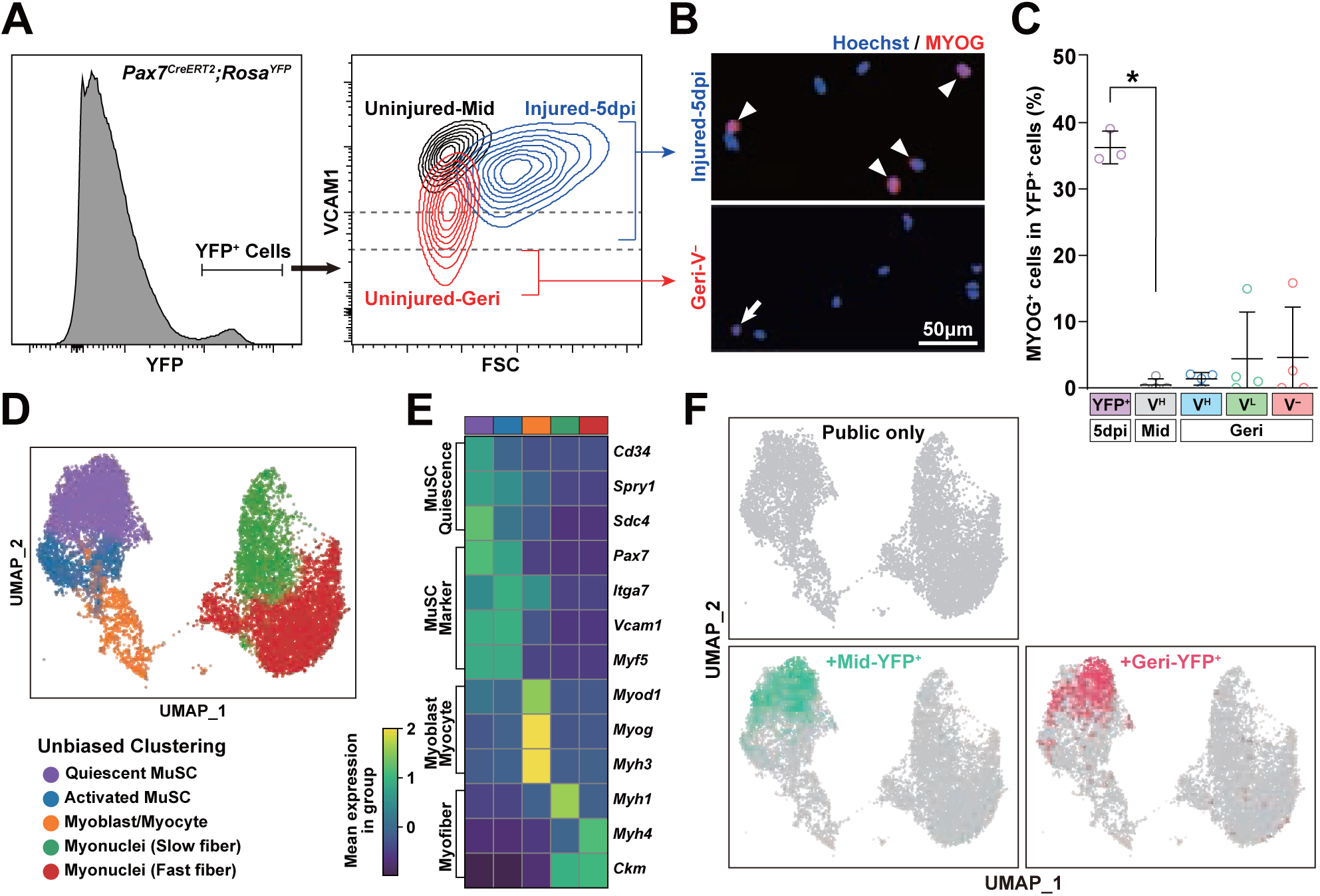
YFP^+^/VCAM1^−^ cells are quiescent MuSCs. (**A**) Representative FACS plot for YFP^+^ cells from uninjured Mid and Geri, and injured-5dpi young (20-weeks-old) mice. (**B**, **C**) Representative images (**B**) of ICC staining for Hoechst (Blue) and MYOG (Red), and quantification (**C**) of MYOG^+^ cells in each YFP^+^ cell, one-way ANOVA followed by Bonferroni’s multiple comparison test; *p* = 0.0139 between YFP+ cells of injured-5dpi and Mid V^H^. While 5 dpi MuSCs differed significantly from young MuSCs (adjusted p = 0.0139), the comparisons between 5 dpi and each geriatric subgroup (VCAM-high, -mid, and -low) did not reach statistical significance after correction for multiple testing (adjusted p = 0.17, 0.15, and 0.17, respectively). (**D**) Uniform manifold approximation and projection (UMAP) plot of YFP^+^ cells integrated with public scRNA-seq data of MuSC-lineage mononucleated cells in uninjured/injured muscle colored by cell types. (**E**) Matrix plot for representative genes of specific cell types for determining the identities of each cluster. (**F**) The localization of Mid and Geri YFP^+^ cells in this study on UMAP plot of Fig. 2D.

To further investigate cell identity, we performed single-cell transcriptome analysis of YFP^+^ cells from middle-aged and geriatric mice and integrated these data into public single-cell sequencing data of uninjured and injured muscles.(De Micheli et al., 2020, Oprescu et al., 2020) (Fig. S1A-E). Integration of public single-cell sequencing data yielded 5 clusters: quiescent MuSC, activated MuSC, myoblast/myocyte, fast fiber myonuclei and slow fiber myonuclei (Fig. 2D). The transcriptional signatures characterizing each cluster were consistent with known lineage identities, supporting the validity of the clustering (Fig. 2E). High levels of myonuclei transcripts likely represents fragmented myonuclei or cells with compromised nuclear integrity, which is a known technical byproduct of dissociation of the myofiber (Santos et al., 2021). Only high-confidence MuSC clusters were included in differential expression, trajectory inference, and cluster-proportion analyses. All YFP^+^ cells from both middle and geriatric age mapped to transcriptional clusters associated with quiescent MuSCs (Fig. 2F). This suggests that even VCAM1^Low^ and VCAM1^−^ cells from geriatric age did not cluster with activated MuSCs or differentiating myoblasts or myocytes, but rather remained within the quiescent MuSC compartment. These findings support the possibility that these cells may retain core stem-like features, despite altered surface marker profiles.

Since quiescent MuSCs are generally smaller than activated progenitors due to reduced cytoplasmic volume, low metabolic activity, and compact chromatin organization(Liu et al., 2013), we also compared the cell size of VCAM1^−^ cells with activated MuSCs from injured mice. Indeed, both VCAM1^Low^ and VCAM1^−^ cells from geriatric mice were similar in diameter to YFP^+^ cells from uninjured middle-aged mice, but were significantly smaller than activated MuSCs from injured mice (Fig. S2A-B). These results are consistent with our interpretation that VCAM1^Low^ and VCAM1^−^ cells from geriatric mice may remain in a quiescent state.

Given reports of smooth muscle mesenchymal cells (SMMCs) expressing overlapping surface markers with MuSCs, we considered whether the VCAM1^−^ cells might reflect SMMC contamination(Giordani et al., 2019). mRNA sequence analysis revealed that this population exhibited low expression of SMMC markers, including Myl9, Acta2, and Pdgfrb (Fig S2C), suggesting they are transcriptionally distinct from SMMCs. VCAM1^−^ cells showed significantly higher Pax7 expression compared to SMMCs. Together, these results suggest that YFP^+^/VCAM1^−^ cells from geriatric age may retain quiescent features and myogenic identity, despite lacking typical MuSC markers.

### Impaired functionality of YFP^+^/VCAM1^−^ MuSCs

Since YFP^+^/VCAM1^−^ cells retained molecular and morphological features consistent with quiescent MuSCs, we next investigated their functional potential. Colony-forming assays revealed that YFP^+^/VCAM1^High^ cells from geriatric mice expanded at comparable rates to those from middle-aged mice. In contrast, colony formation by YFP^+^/VCAM1^Low^ and YFP^+^/VCAM1^−^ subsets were progressively reduced, with the VCAM1^−^ subsets forming smaller colonies (Fig. 3A-B). These findings suggest that these cells retain proliferative ability, although their expansion capacity was compromised.

**Figure 3.**
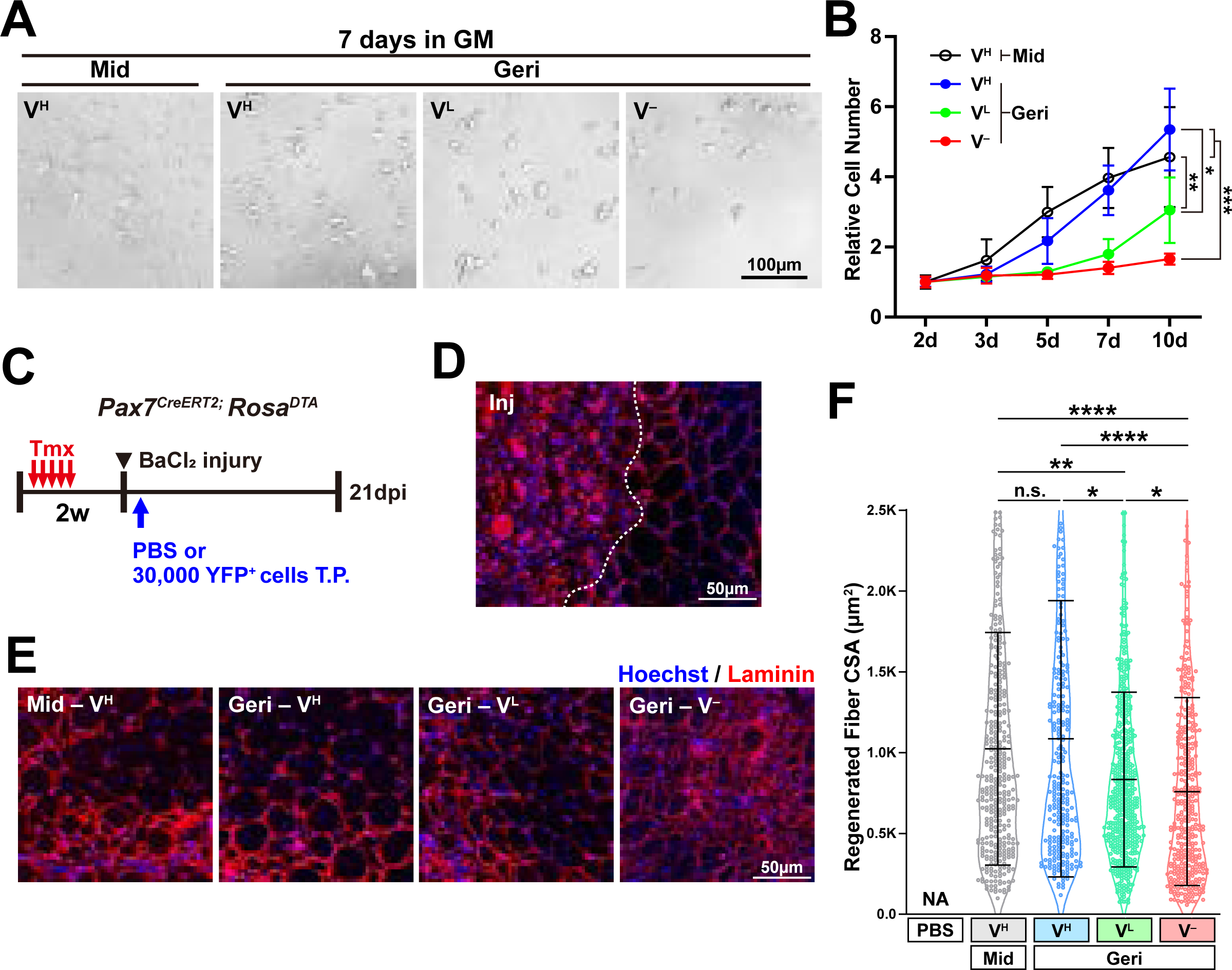
Compromised but retained functionality in VCAM1^−^ MuSCs. (**A**) Representative bright field images of Mid V^H^, Geri V^H^, V^L^, and V^−^ MuSCs at 7 days culture in MuSC growth media. (**B**) Proliferation measured by relative cell number at 2, 3, 5, 7, and 10 days, two-way ANOVA followed by Bonferroni’s multiple comparison test; *p* = 0.0029 and 0.0002 (Mid V^H^ vs V^L^ and V^−^, respectively) and *p*= 0.0102 and 0.0006 (Geri V^H^ vs VL and V^−^, respectively). For each sample, 30,000 MuSCs were plated at day 0, with 3 biological replicates per condition. (**C**) Experimental schematics of transplantation of YFP^+^ cells to TA muscle of *Pax7^CreERT2^;Rosa^DTA^*mice. (**D, E**) Representative images of IHC staining for Hoechst (Blue) and Laminin (Red) in PBS treated TA muscle section following injury. (**D**) Left side of the dotted line show areas affected by BaCl2 injury and right side shows uninjured areas of the muscle. (**E**) IHC staining shows centralized myofibers in all images, with notably smaller myofibers in Geri V^−^. (**F**) Quantification of cross-sectional area (CSA) of regenerated fibers from TA muscle of *Pax7^CreERT2^;Rosa^DTA^*mice transplanted with Mid V^H^, Geri V^H^, V^L^, and V^−^ at 21 dpi, n = 3 for each group. For each mouse, the entire TA muscle was cryosectioned and immunostained, and all regenerated fibers containing centrally located nuclei were included in the CSA quantification. Total 43 data points are outside of the axis. Kruskal-Wallis test followed by Dunn’s multiple comparison test; p < 0.0001 (Mid and Geri V^H^ vs V^−^) and *p* = 0.0059 (Mid V^H^ vs Geri V^L^), 0.0149 (Geri V^H^ vs V^L^) and 0.0143 (Geri V^L^ vs V^−^).

To examine their regenerative potential *in vivo*, we transplanted equal numbers of YFP^+^ cells from each group into pre-injured TA of MuSC-deficient *Pax7^CreERT2^;Rosa^DTA^*recipient mice (Fig. 3C). In the absence of donor cells, regeneration was severely impaired, consistent with a lack of endogenous MuSCs (Fig 3D). Transplantation of YFP^+^/VCAM1^High^ cells from both middle-aged and geriatric donors supported robust regeneration, resulting in centrally nucleated myofibers with comparable morphology between age groups, consistent with heterotypic transplantation studies (Fig. 3D-E) (Ikemoto-Uezumi et al., 2015). Interestingly, both YFP^+^/VCAM1^Low^ or YFP^+^/VCAM1^−^ cells were also capable of contributing to myofiber regeneration, despite lacking traditional MuSC markers (Fig. 3E). However, regenerated myofibers in these groups were significantly smaller than those derived from VCAM1^High^ cells (Fig. 3F), suggesting that loss of VCAM1 expression may be associated with diminished regenerative capacity.

Together, these results indicate that even the most marker-deficient MuSCs in geriatric muscle, YFP^+^/VCAM1^−^ cells, retain a degree of regenerative competence. However, their reduced proliferative output and attenuated *in vivo* contribution suggests functional heterogeneity within the aged MuSC pool, wherein diminished marker expression may correlate with impaired regenerative performance.

### VCAM1^−^ MuSCs exhibit senescence-like features and sensitivity to senolytics

To investigate the molecular basis of impaired function in VCAM1^−^ MuSCs, we performed mRNA sequence analysis on YFP^+^ MuSCs. Samples showed high quality and a strong correlation was observed among biological replicates (Fig. S3A-C). Each group consistently reproduced similar gene expression patterns in biological replicates in hierarchical clustering (Fig. S3D). Although mid-aged VCAM1-high MuSCs did not exhibit the emergence of a VCAM1-low state, mRNA profiling revealed detectable transcriptional differences between mid-aged and geriatric VCAM1-high populations (Fig. S3D). These shifts likely reflect the gradual accumulation of aging-associated molecular changes that begin during the middle-aged stage, even before overt functional decline or loss of VCAM1 becomes evident. Importantly, these early transcriptomic alterations do not give rise to the dysfunctional VCAM1-low subset, which emerges only in geriatric muscle. In particular, hierarchical clustering showed similar gene expression pattern between VCAM1^High^ MuSCs of middle-aged and geriatric mice and between VCAM1^Low^ and VCAM1^−^ MuSCs (Fig. S3E), consistent with our results of *in vivo* and *in vitro* functionality experiments (Fig. 3A-F).

Kyoto-encyclopedia of genes and genomes (KEGG) pathway enrichment analysis based on the DEGs (Fig. S3E) (Kolberg et al., 2023) showed alterations in pathways related to cellular senescence and PI3K-AKT pathway in VCAM1^−^ MuSCs of geriatric age as compared to VCAM1^High^ MuSCs of middle-age (Fig. 4A). We further investigated the expression levels of individual genes known to be altered in MuSCs of aged mice, such as genes related to extracellular matrix (Schuler et al., 2021), Notch signaling (Liu et al., 2018, Conboy et al., 2003, Conboy et al., 2005), and autophagy-lysosomal pathway (Garcia-Prat et al., 2016, Kim et al., 2021b), some of which were more altered in VCAM1^Low^ and VCAM1^−^ MuSCs compared to VCAM1^High^ MuSCs of geriatric mice (Fig. 4B). *Cdkn2a* (*p16^INK4a^*) and *Cdkn2b* (*p15^INK4b^*), which are cell cycle regulators widely used as markers for senescent cells, showed highly altered expression patterns in VCAM1^−^ MuSCs compared to VCAM1^Low^ MuSCs (Fig. 4B) (Saito and Chikenji, 2021). However, the expression levels of other well-known cellular senescence marker genes, such as *Rb1*, *Il6*, and *Trp53*, were not altered much. *Pik3cb* and *Tsc1,* which are involved in the PI3K-Akt-mTOR pathway, showed overall increased expression, while other genes in the PI3K-Akt pathway remained relatively unchanged. Together, these results suggest a unique combination of *in vivo* senescence-like molecular characteristics of VCAM1^−^ MuSCs, distinct from molecular characteristics of replicative cellular senescence.

**Figure 4.**
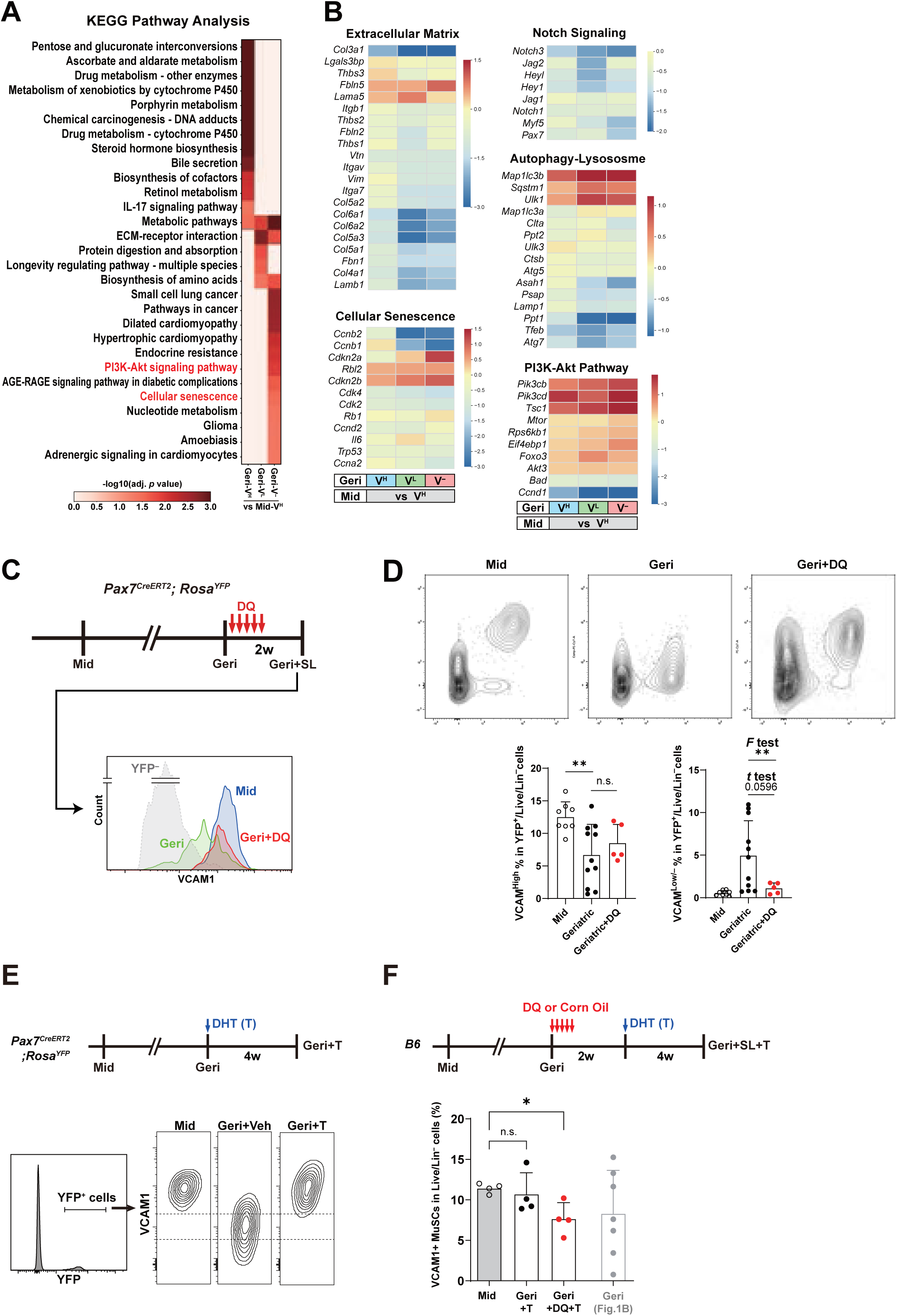
Senescence characteristics of VCAM1^−^/YFP^+^ cells and their rejuvenation by DHT supplementation. (**A**) The statistical significance of Kyoto-Encyclopedia of Genes and Genomes (KEGG) pathways, associated with differentially expressed genes (DEGs) between Mid V^H^ and Geri V^H^, V^L^ and V^−^, visualized as a heatmap. (**B**) Heatmap of relative expressions of representative genes known to be altered in aged MuSCs compared to Mid V^H^. (**C**) Experimental schematics of senolytics, dasatinib and quercetin (DQ), treatment in MuSC^YFP^ at geriatric age and representative FACS histogram of VCAM1 expression Mid, Geri, and Geri treated with SL (Geri+DQ) mice. (**D**) Representative FACS plots and quantification of the percentage of VCAM1^High^ and VCAM1^Low/^**^−^** cells divided by VCAM1 expression levels in Mid (n = 8) and Geri mice with (n = 5) or without (n = 11) senolytics, 2 tailed student’s t test, *p* = 0.0051 for Mid vs Geri in VCAM1^High^ and *F* test: *p* = 0.0026 for Geriatric vs Geriatric+DQ in VCAM1^Low/–^. (**E**) Experimental schematics of dihydrotestosterone (DHT) treatment and FACS plot of VCAM1 expression levels for YFP^+^ cells in MuSC^YFP^. (**F**) Experimental schematics of DQ and DHT treatment and quantification of the percentage of VCAM1^High^ MuSCs in Live/Lin– cells in Mid and Geri B6 mice (n = 4 in each), 2 tailed student’s t test, *p* = 0.0113 for Young vs Geri+DQ+T.

Given their unique senescence associated transcriptome and partially elevated PI3K-Akt pathway, we tested whether VCAM1^−^ MuSCs were susceptible to senolytic elimination. Geriatric mice were treated with senolytic agents dasatinib and quercetin (DQ) (Xu et al., 2018), and analyzed 2 weeks later (Fig. 4C). Despite variation in baseline VCAM1^High^ and VCAM1^Low/–^ population across aged animals, a consistent inverse correlation was observed between VCAM1^High^ and VCAM1^Low/–^ fractions (Fig. S3F). The multimodal distribution observed in the geriatric group reflects increased mouse-to-mouse variability that emerges at very advanced ages. Following DQ treatment, VCAM1^High^ MuSCs remained comparable while VCAM1^Low/–^ MuSCs showed reduced abundance and lower variability (p = 0.0596) (Fig. 4D). The multimodal distribution observed in the geriatric group reflects increased mouse-to-mouse variability that emerges at very advanced ages. Although aging affects all animals, the extent and timing of phenotypic changes vary substantially among individuals, resulting in broader divergence in VCAM1 expression states. Statistical analyses were performed using the mouse as the biological replicate, this variability does not alter the overall conclusion. These results indicate selective sensitivity of VCAM1^Low/–^ MuSCs to senolytic agents and their increased dependence on BCL-2–related survival pathways.

Together, our findings reveal a distinct subset of quiescent MuSCs in geriatric muscle that exhibit unique senescence-like molecular features and are amenable to senolytic clearance. We term these cells GERI-MuSCs (Geriatric Emerging Regeneration-capable and Impaired-MuSCs).

### Rejuvenation of GERI-MuSCs by DHT treatment

Studies have demonstrated that treatment with rejuvenating agents can restores myofiber regeneration capacity in aged mice (Bernet et al., 2014, Cosgrove et al., 2014, Zhang et al., 2016, Elabd et al., 2014, Price et al., 2014, Garcia-Prat et al., 2016). Previously, dihydrotestosterone (DHT) supplementation has been shown to preserve MuSCs function and enhance regeneration in Antide, a GnRH antagonist, treated mice by restoring autophagy activation (Kim et al., 2021b). Consistent with these findings, implantation of DHT-filled silastic tube in 28-month-old geriatric mice improved muscle regeneration, as evidenced by increased fiber cross sectional area (Fig. S4A-C).

To investigate whether DHT could affect GERI-MuSCs, we treated 28-month-old MuSC^YFP^ mice with DHT and analyzed MuSC populations one month later (Fig. 4E). Remarkably, DHT treatment resulted in a decrease in GERI-MuSCs with a corresponding expansion of VCAM1^High^ MuSCs population. This phenotypic reversion suggests that DHT promotes restoration of canonical marker expression in GERI-MuSCs.

To determine whether DHT acts through rejuvenation of GERI-MuSCs, we first treated the mice with DQ for 5 consecutive days and then implanted DHT tube 2 weeks later (Fig. 4F). In mice pre-treated with DQ, DHT supplementation failed to reconstitute the VCAM1^High^ MuSC pool (Fig. 4F). This finding suggests a model in which GERI-MuSCs are not terminally senescent, but rather exist in a latent, hormone-responsive state. Upon DHT exposure, these cells likely reacquire a VCAM1^High^ phenotype, underscoring their potential as a therapeutic target in muscle aging.

### Identification of GERI-MuSCs using CD63 and CD200

In non-reporter models, the detection and isolation of GERI-MuSCs pose a significant challenge, as these cells lack expression of conventional MuSC markers such as VCAM1 and PAX7 at geriatric age. To overcome this limitation, we conducted a transcriptomic screen to identify alternative surface markers that remain stably expressed across the full spectrum of MuSC aging states. Candidate genes were selected based on three criteria; high transcript abundance (RPKM), consistent expression from VCAM1^High^ to VCAM1^−^ MuSCs, and predicted surface localization (Fig. 5A). CD63 and CD200 sufficed all three criteria and were therefore chosen as candidate markers for further validation (Fig. 5B).

**Figure 5.**
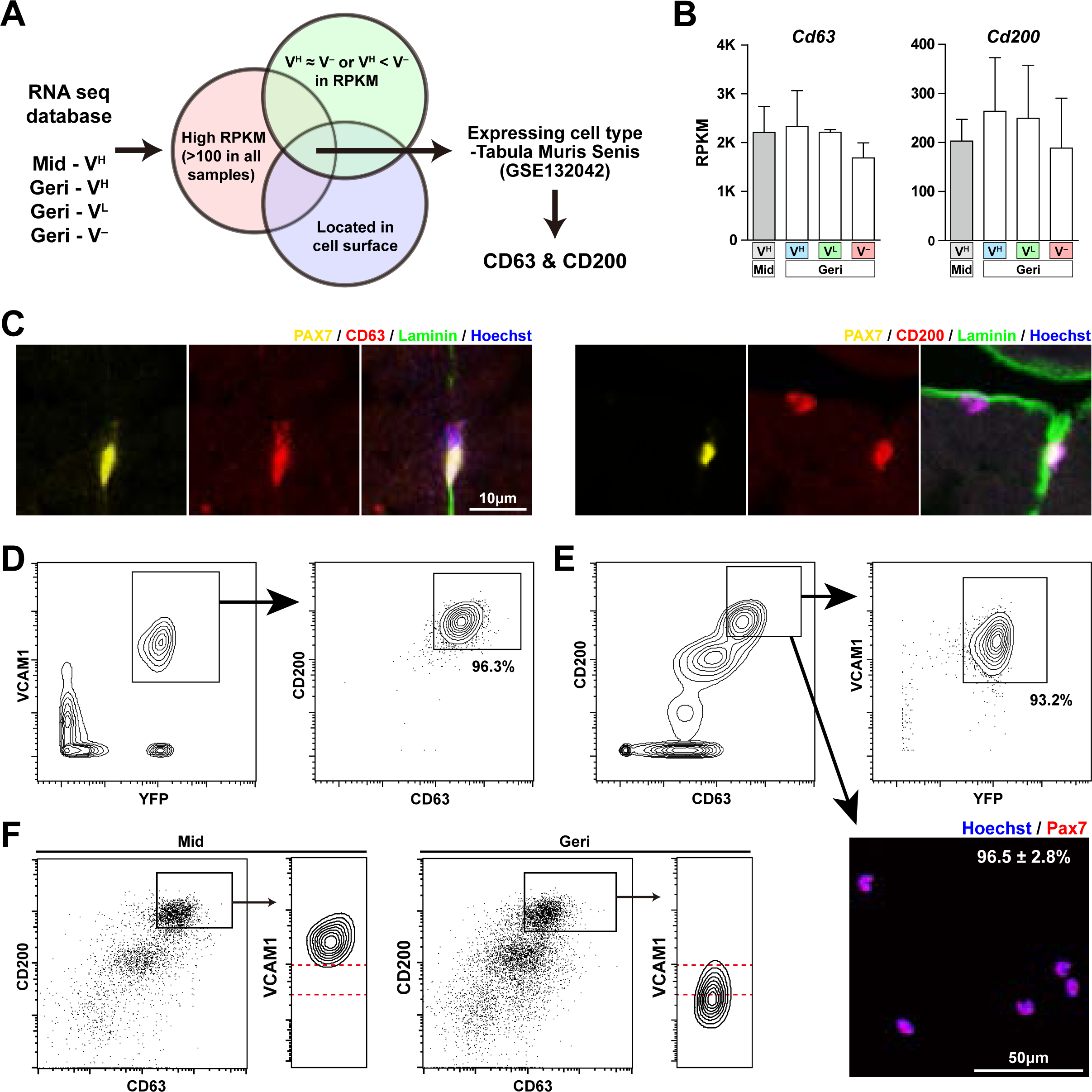
Detection of GERI-MuSCs using CD63/CD200-based FACS strategy. (**A**) Vann Diagram schematics showing criteria used to select MuSC marker candidate CD63 and CD200, from RNA seq database. (**B**) Relative gene expressions of *Cd63* and *Cd200* shown in RPKMs from Mid V^H^, Geri V^H^, V^L^, and V^−^ MuSCs. (**C**, **D**) Representative IHC staining images for Hoechst (Blue), Laminin (Green), PAX7 (Yellow) with CD63 (red, **C**) or CD200 (red, **D**) in the TA muscle of Mid MuSC^YFP^. (**E**) Representative FACS plot from young mice showing 96.3% of VCAM1^+^/YFP^+^ cells are CD63^high^/CD200^high^. (**F**) Left: representative FACS plot from young mice showing 93.2% of CD63^high^/CD200^high^ cells are VCAM1^+^/YFP^+^, and ICC staining of this population shows that 96.5±2.8% are PAX7^+^. Right: representative FACS plot of MuSCs of *B6* Mid and Geri mice gated by CD63 and CD200 and their VCAM1 expression level.

We confirmed protein level expression of CD63 and CD200 expression in MuSCs by co-immunostaining TA cross-sections for CD63, CD200, YFP and PAX7. Both markers co-localized with YFP and PAX7 beneath the basal lamina (Fig. 5C). Flow cytometry analysis further demonstrated that 96.3% of YFP^+^/VCAM1^+^ cells from young MuSC^YFP^ mice were CD63^high^/CD200^high^ (Fig. 5D). Notably, among lineage-negative (CD31^−^/CD45^−^), SCA-1^−^, and live cells, 93.2% of CD63^high^/CD200^high^ cells were YFP^+^/VCAM1^+^ cells (Fig. 5E). Importantly, 96.5±2.8% of sorted CD63^high^/CD200^high^ cells were PAX7^+^, validating these markers as a robust strategy for MuSC enrichment.

To assess whether the CD63/CD200-based strategy could detect and isolate GERI-MuSCs in wild-type mice, we applied the strategy to middle and geriatric aged mice. In middle aged mice, all CD63^high^/CD200^high^ cells corresponded exclusively to VCAM1^High^ MuSCs (Fig. 5F). In contrast, geriatric mice exhibited a marked shift, with the majority of CD63^high^/CD200^high^ cells falling within the VCAM1^Low/–^ gate (Fig. 5F). These results indicate that CD63 and CD200 expression is preserved in GERI-MuSCs, enabling their prospective identification and isolation in the absence of genetic labeling. Therefore, these findings establish CD63 and CD200 as reliable surface markers that can be used with VCAM1. Together, they can to detect the full spectrum of MuSCs, including functionally latent populations that evade detection by conventional means. This strategy offers a critical tool for future studies aiming to comprehensively characterize MuSC heterogeneity and therapeutic responsiveness in the context of aging.

## Discussion

Our study challenges the longstanding paradigm that muscle stem cell (MuSC) number declines irreversibly with age. While previous reports have described significant depletion of Pax7^+^ MuSCs in aged muscle, these findings were largely based on immunohistochemical detection or flow cytometric profiling using conventional surface markers. Here, we demonstrate that genetically labeled MuSCs, tracked longitudinally *in vivo* using a Pax7-CreERT2;Rosa-YFP model, do not decrease in number even in geriatric mice. Instead, a substantial subset of aged MuSCs loses expression of canonical markers such as Pax7 and VCAM1, rendering them undetectable by traditional approaches. Although VCAM1 is frequently used as a surface marker for MuSC identification, it represents only one of several commonly used markers, and ITGA7-based strategies remain widely applied in the field. In our dataset, ITGA7 expression within the YFP⁺ MuSC population did not show a significant reduction in mean fluorescence with aging (Fig. 1D). However, the variance of ITGA7 expression was markedly increased in geriatric MuSCs, indicating greater phenotypic heterogeneity. This pattern suggests that aging does not uniformly downregulate canonical markers but instead increases instability and variability in their expression, which may lead to under-detection of a subset of MuSCs even when ITGA7 is used as the primary gating marker. These findings underscore that marker-based quantification may underestimate specific aging-associated MuSC subpopulations and should be interpreted within this limitation.

We identify this overlooked population as GERI-MuSCs (Geriatric Emerging Regeneration-capable and Impaired MuSCs). Despite their diminished expression of hallmark MuSC markers, GERI-MuSCs exhibit features consistent with quiescence, rather than terminal differentiation or apoptosis. Although GERI-MuSCs showed markedly reduced colony formation and further validation using direct viability assays are necessary, several lines of evidence suggest that they are viable rather than acutely undergoing cell death. These cells can be isolated as intact populations by FACS and generate high-quality RNA-sequencing libraries, which would not be feasible if they were non-viable. Their transcriptomic profiles also show enrichment of stress-response and senescence-associated programs rather than apoptotic signatures. Together, these findings suggest that GERI-MuSCs remain alive but exhibit impaired proliferative capacity. Furthermore, transcriptomic profiling and functional assays reveal that these cells retain a partial capacity for proliferation and regeneration, albeit at reduced efficiency. When comparing geriatric VCAM1^+^ MuSCs with middle aged MuSCs, we found 1,428 DEGs, where 701 genes were downregulated and 727 genes were upregulated (Fig. S3E). Some of the pathways altered were similar to previously reported differences, such as alterations in the autophagy-lysosome related genes and PI3K-Akt Pathways (Garcia-Prat et al., 2016). However, these transcriptomic alterations did not affect the functional integrity of geriatric VCAM1^+^ MuSCs (Fig. 3 A-F). Taken together, these results do not contradict prior studies demonstrating age-related MuSC decline but rather refine this understanding by highlighting a subset of aged MuSCs that may have been underrepresented or more difficult to detect using conventional markers. By resolving these cells with an alternative phenotypic definition, our findings complement existing models of MuSC aging and suggest that part of the apparent depletion previously reported may reflect changes in marker expression rather than complete cellular loss.

GERI-MuSCs display a unique gene expression profile that includes upregulation of senescence-associated transcripts such as *Cdkn2a* but unaltered expression of other typical senescence-associated genes, such as *Il6* and *Trp53*, while showing preservation of certain stemness-related pathways (Fig. 4B). These cells do not fulfill the classical definition of cellular senescence, such as irreversible cell cycle arrest(Campisi and d’Adda di Fagagna, 2007). p16 expression is elevated in VCAM-low MuSCs, but this alone does not establish a fully senescent state. Rather, GERI-MuSCs exhibit a hybrid state of retaining regenerative potential while expressing select markers traditionally associated with senescence. This finding suggests that the criteria used to define senescence in proliferative cell types may not fully apply to quiescent stem cells like MuSCs, which spend most of their life in a non-proliferative state. Additional markers and functional assays will be needed to define the precise state of these cells and to delineate *in vivo* senescence features specific to long-lived, tissue-resident stem cells.

Our results suggest that GERI-MuSCs could be functionally reactivated. Treatment with dihydrotestosterone (DHT) restored VCAM1 expression in GERI-MuSCs and regenerative performance in aged mice, suggesting that their phenotype is reversible and hormonally responsive. This provides a compelling explanation for the efficacy of hormone-based interventions in aging muscle and highlights the potential for targeting latent regenerative capacity in aged stem cells. Notch signaling appears to be a key regulator of this reversible phenotype. We found diminished expression of Notch pathway components, including Notch1, Notch3, Hey1, and HeyL, in VCAM1^Low^ and VCAM1^−^ subsets, along with reduced transcription of Pax7 and VCAM1. Given that both Pax7 and VCAM1 are downstream targets of Notch signaling and that Notch activity can be modulated by androgen signaling, the restoration of MuSC function by DHT may act through this axis. Testing such mechanisms in future studies will be important for establishing therapeutic strategies.

Although DHT supplementation partially restored the regenerative performance of geriatric MuSCs, this effect cannot be attributed solely to the rescue of VCAM1 expression in GERI-MuSCs. Androgen signaling is known to influence multiple non–stem cell populations in the muscle environment, including fibro-adipogenic progenitors, myofibers, and immune cells, any of which may contribute to the observed improvement in regeneration (Kim et al., 2021b). Accordingly, we acknowledge that DHT likely exerts broader tissue-level effects beyond the MuSC compartment, and our findings should be interpreted within this limitation. Future studies combining senolytic strategies with targeted modulation of androgen signaling may help delineate the relative contributions of each pathway, although such experiments extend beyond the scope of the present work.

The presence of GERI-MuSCs also raises caution about the use of senolytic agents. Although dasatinib and quercetin are used to clear senescent cells and have shown benefits in aged tissue, our data show that these agents also deplete GERI-MuSCs. Since GERI-MuSCs retain functionality and regenerative potential, their removal could compromise muscle maintenance and repair in elderly individuals. Given the critical role of MuSCs in preserving muscle mass, supporting myonuclei turnover, and maintaining neuromuscular junctions, the long-term application of senolytics in skeletal muscle warrants careful scrutiny.

Finally, we propose CD63 and CD200 as reliable surface markers that can be used with VCAM1 for identifying MuSCs across age groups. These markers remain expressed in both young and aged MuSCs, including those with diminished canonical marker expression. CD63 and CD200 expression in geriatric muscle was evaluated using conventionally isolated MuSCs rather than YFP-traced cells. This distinction should be considered when interpreting these results. Their use enables comprehensive isolation of the full MuSC population and will be instrumental in improving the resolution of future studies on stem cell heterogeneity and aging.

In conclusion, we identify a previously unrecognized population of MuSCs—GERI-MuSCs—that persists with age despite loss of conventional markers. These cells remain quiescent, partially functional, and responsive to rejuvenation cues. Their discovery challenges existing models of MuSC depletion and senescence, and underscores the importance of revisiting stem cell aging with improved detection strategies and nuanced interpretations of cellular function.

## Materials and Methods

### List of Antibodies for Flow Cytometry and FACS (Fluorescence-activated cell sorting

*List of Antibodies for Flow Cytometry and FACS (Fluorescence-activated cell sorting)*
| Marker | Fluorophore | Catalog No. | Company |
| --- | --- | --- | --- |
| CD31 | APC | 102510 | BioLegend |
| CD31 | Pacific Blue™ | 108120 | BioLegend |
| CD45 | APC | 103112 | BioLegend |
| CD45 | Pacific Blue™ | 405206 | BioLegend |
| VCAM1 | Biotin | 553331 | BD Biosciences |
| Streptavidin | PE/Cy7 | 405206 | BioLegend |
| Ly-6A/E (Sca-1) | Pacific Blue™ | 108120 | BioLegend |
| Ly-6A/E (Sca-1) | FITC | 553335 | BD Biosciences |
| CD200 | BV711 | 745548 | BD Biosciences |
| CD63 | APC | 170631-82 | eBioscience |

### Animals

All animal procedures conformed to the guidelines of Institutional Animal Care and Use Committee (IACUC) of Seoul National University, Korea. Male C57BL/6J mice were housed under a 12-h light/12-h dark cycle at 22°C with 40-60% humidity, and fed ad libitum. Pax7^CreER^ (017763), Rosa^YFP^ (006148), Rosa^DTA^ (009669) mice were obtained from the Jackson Laboratory (Bar, Harbor, ME, USA). Tamoxifen (25mg/mL in corn oil; Sigma-Aldrich, St. Louis, MO, USA) was administered orally for 5 consecutive days (240 mg/kg). For muscle injury, mice were anesthetized with avertin (250 μg/g, i.p.) and injected with 60 μL of 1.2% BaCl2 (Sigma-Aldrich) into the Tibialis Anterior (TA) muscles. One-day post-injury, mice were anesthetized with avertin and MuSCs were transplanted at indicated numbers.

### Flow Cytometry and FACS (Fluorescence-activated cell sorting)

Limb muscles were mechanically dissected and enzymatically dissociated in Dulbecco’s modified Eagle’s medium (DMEM) containing 10% horse serum (Hyclone, Logan, UT, USA), collagenase type II (800U/mL; Worthington, Lakewood, NJ, USA), and dispase (1.1 U/mL; StemCell Technologies, Vancouver, BC, Canada) at 37°C for 50 minutes. Digested suspensions were triturated, washed with DMEM and filtered to obtain mononuclear cells. Cells were stained with antibodies, analyzed and sorted using a FACS Aria III or FACS Aria Fusion (BD Bioscience). List of antibodies used are provided as supplementary materials. For all flow cytometry analyses, fibro-adipogenic progenitors (FAPs) were excluded by gating out Sca1⁺CD31⁻CD45⁻ cells prior to MuSC identification. This strategy ensured that YFP⁺ MuSC quantification was not confounded by age-associated changes in Sca1⁺ mesenchymal populations. Representative plots were processed with FlowJo v10.9.

### MuSC Culture

FACS-isolated MuSCs were cultured in DMEM/F12 + 20% FBS + 1% ZellShield (Minerva Biolabs, Germany) + 2.5 ng/mL bFGF (F0291, Sigma Aldrich) on Matrigel (256231, BD Bioscience)-coated plates at 37°C and 5% CO2. Medium was refreshed every 3 days. MuSCs were imaged daily using EVOS FL Auto 2 (Thermo Fisher, USA) and the number of cells in the same region was counted.

### Histochemistry and Immunohistochemistry

TA muscles were fixed overnight in 4% paraformaldehyde, embedded in the Tissue Tek OCT compound (Sakura Fineteck, USA), and snap-frozen in liquid nitrogen. The muscles were sectioned at 7μm and stained on the slides. H&E staining was performed according to basic protocols. For immunostaining, sections were subjected to epitope retrieval in 50mM Tris-EDTA (pH 9.0) at 105°C for 10 minutes, then blocked with 3% BSA + 0.1% Triton X-100. Sections were incubated overnight with primary antibodies, mouse anti-PAX7 (DSHB, Pax7, 1:100 dilution), chicken anti-GFP (abcam, ab13970, 1:300), rabbit anti-laminin (Sigma, L9393-2ML, 1:500), rat anti-CD200 (BD Biosciences, 1:50), and rat anti-CD63 (eBioscience, 1:50). After washing, sections were incubated with secondary antibodies, Alexa Fluor 647-conjugated anti-rat or anti rabbit IgG, Alexa Fluor 594-conjugated anti mouse IgG, and Alexa Fluor 488-conjugated anti chicken IgG. Nuclei were counterstained with Hoechst 33342 (Invitrogen, H3570). Slides were mounted with VECTASHIELD (Vector Laboratories) and images were acquired using EVOS FL Auto 2 (Thermo Fisher) and Leica TCS SP8 confocal microscope. Cross-sectional area (CSA) was calculated by SMASH[23] and ImageJ programs, respectively.

### Immunocytochemistry

Isolated MuSCs were cytospun onto slides (Cytospin 4, Thermo Scientific), fixed with 4% paraformaldehyde for 10 minutes, washed with PBS and permeabilized with PBS containing 0.4% Triton X-100 for 30 minutes. Slides were then incubated with primary antibodies, mouse anti-PAX7 (DSHB) and mouse anti-MYOG (DSHB, F5D, 1:100 dilution) in blocking buffer. Subsequent steps followed the immunohistochemistry protocol. Relative fluorescent intensity was quantified using ImageJ.

### Senolytics and DHT treatment

Dihydrotestosterone (DHT) implantation was performed as previously described[8]. Senolytic agents Dasatinib (D-3307, LC laboratories, 5mg/kg) and Quercetin (Q4951, Sigma, 50 mg/kg) was administered orally for 5 consecutive days. Experimental schemes of combined treatment are as mentioned in the results section of this article.

### Single-cell RNA sequence analysis

To obtain gene-specific UMI (Unique Molecular Identifier) counts in each cell, we processed raw scRNA-seq data using the BD Rhapsody WTA analysis pipeline (v1.11.1)[24]. Scanpy (v1.9.3)[25] was utilized for single-cell RNA sequence analysis. Cells with a gene count below 200 or a mitochondrial gene percentage above 20% were considered low-quality cells and were excluded from downstream analyses. We set the upper limit of each UMI count as (mean(total counts) + 2 standard deviation(total counts) value) to remove doublets. Normalization was applied to the count matrix (scanpy.pp.normalize_total with target_sum=10^4^ and scanpy.pp.log1p).

We characterized the enriched cells by merging our scRNA-seq data with public datasets (GSE162172, GSE138826)[26,27]. From the public dataset, only the cells belonging to the myogenic lineage (from quiescent MuSCs to Myonuclei (Type IIb or IIx)) were selected for comparison. After merging the public dataset with our dataset (scanpy.external.pp.harmony_integrate with max_iter_harmony=50), 2000 highly variable genes were identified and scaled (scanpy.pp.highly_variable_genes with n_top_genes=2000, scanpy.pp.scale with max_value=10).

We visualized the scRNA-seq data using uniform manifold approximation and projection (UMAP) (scanpy.tl.umap) with a precomputed neighborhood graph (scanpy.pp.neighbors with n_pcs=50). We clustered the UMAP-transformed data using the Louvain algorithm (scanpy.tl.louvain, resolution=0.5). Each cluster was annotated using cell-specific markers (Fig. 1D, E). The proportions of quiescent MuSCs, activated MuSCs, and Myoblasts/Progenitors were measured in our samples (Mid and Geri) and the public dataset across various days-post-injury (DPI). Cases with less than 50 cells per DPI were excluded during this analysis (Fig. S1A).

We utilized diffusion map, PAGA, and force-directed graph to analyze the trajectory of cells belonging to the myogenic lineage. After dimension reduction using diffusion maps (scanpy.tl.diffmap), we computed neighborhood graphs (scanpy.pp.neighbors) and abstracted the graphs using PAGA (scanpy.tl.paga). We modeled the trajectory of the cells using the abstracted graphs as initial positions by constructing a force-directed graph (Fig. S1B-E).

### Representative Isoforms of Mouse mRNAs

To acquire a sole representative isoform for each gene, we utilized the filtering pipeline described in a previous study(Kim et al., 2016). The mouse reference annotation (GRCm39 refFlat) was downloaded from the UCSC website(Pruitt et al., 2007). mRNA sequences with accession category ‘NM’ located on the 21 chromosomes (1-19, X, and Y) were selected for downstream analyses. Entries with incorrect start or stop codons, internal stop codons, and ORF lengths not divisible by 3, were removed. Nonsense-mediated mRNA decay candidates, defined as the mRNAs with stop codons positioned less than 50 nucleotides before the final exon-exon junction, were excluded (Nagy and Maquat, 1998). A total of 20,582 non-redundant representative isoforms were selected as mouse reference mRNAs for subsequent analyses.

### mRNA sequence analysis

mRNA sequence analysis was performed on 3-4 biological replicates for each condition. We trimmed low-quality read ends using a custom algorithm. Each base position was assigned a reward (+5) or a penalty (-1) based on its Phred score with a threshold of 30. To define high-quality read ends, we calculated the cumulative sum from both ends, identified the positions with the minimum value, and trimmed the reads at those positions. This process was repeated iteratively until no further trimming was necessary. Reads shorter than 17 bases after trimming were excluded. Subsequently, we removed adapter sequences using Cutadapt (v3.1) (Martin, 2011) with options (-O 10 -m 18 --match-read-wildcards). We eliminated sequencing artifacts and PCR duplicates using FASTX-Toolkit’s fastx_artifact_filter with option (-Q 33) (hannonlab.cshl.edu/fastx_toolkit/, v0.0.13). To eliminate reads derived from mouse non-coding RNA, we aligned the reads to non-coding RNA sequences in RNAcentral(Consortium, 2021) using Bowtie2 (v2.1.0) (Langmead and Salzberg, 2012) with options (-k 1 --norc --very-sensitive), and the reads aligned to non-coding RNAs were excluded. Unpaired reads were also excluded.

Preprocessed reads were aligned to the mouse genome (GRCm39) utilizing the STAR alignment software (v2.7.5b) (Dobin et al., 2013). The parameters used for the STAR alignment are described in a previous study (Kim et al., 2021a). The reads were sorted and indexed using Samtools (v0.1.19) (Danecek et al., 2021). The expression of each gene was quantified by calculating the fragment per kilobase of transcript per million mapped reads (FPKM).

We applied the same analysis pipeline to both our experimental samples (Mid, V^H^, V^L^, V^-^) and publicly available SMMC samples (GSE110878, SRR67542132-SRR67542134)(Giordani et al., 2019) to calculate gene expression levels. Quantile-normalized expression levels were compared between the two datasets (Fig. S2C).

The following analyses were performed exclusively on our experimental samples.

To assess the quality of our samples, the proportion of reads that were either removed, unmapped, or successfully mapped was computed for each sample throughout the preprocessing and alignment steps (Fig. S3B). The correlation between replicates was quantified using Spearman’s R (Fig. S3C).

We normalized all samples using the following procedure to ensure a fair inter-sample comparison. We applied quantile normalization to equalize the FPKM distribution of all samples. To remove individual mouse variations, we used the RUVs function (k=2) from the RUVSeq package (v1.34.0) (Risso et al., 2014). Subsequently, genes with normalized expression higher than the 50^th^ percentile were selected, and the expression values were z-normalized. We clustered the samples using the clustermap function of the Seaborn package (v0.11.1) (Waskom, 2021) with correlation metric (Fig. S3D).

We identified differentially expressed genes (DEGs) by comparing the expression between the Mid, V^H^, V^L^, and V^-^ samples. We used DESeq2 (v1.40.2) (Love et al., 2014) to compute log^2^ fold changes and false discovery rate (FDR) corrected *Q* values for individual genes. DEGs were defined as genes with |log^2^ fold change| > 1.2 and FDR-corrected *Q* value < 0.05 (Fig. S3E).

To annotate the pathways linked to DEGs, we performed enrichment analysis for Kyoto Encyclopedia of Genes and Genomes (KEGG) pathways using the g:Profiler web server (Kolberg et al., 2023, Kanehisa et al., 2016). Among the highly expressed 12,000 genes, DEGs were queried using the g:GOst tool, and the g:SCS algorithm was employed for multiple testing correction. Pathways with multiple testing corrected *Q* value < 0.05 were presented (Fig. 3A).

### Statistical Analysis

All statistical analyses were performed using R (R-4.3.1) and GraphPad Prism. All variables were confirmed to have a normal distribution by using the Shapiro-Wilk test and Kolmogorov-Smirnov test. For comparison of significant differences in 2 groups of normally distributed data, Levene’s test for homogeneity of variance using the car package followed by a 2-tailed Student’s *t* test or Welch’s *t* test was used. For comparison of multiple groups, one-way analysis of variance (ANOVA) or two-way ANOVA followed by Bonferroni’s multiple comparison test was performed. All data are presented as mean ± s.d. n.s., not significant; P > 0.05; *P <0.05; **P<0.01; ***P<0.001, ****P<0.0001. *P value* of <.05 considered statistically significant at the 95% confidence level. The number of biological replicates, *p* values, and detailed statistical method for each data were indicated in the figure legends.

## Acknowledgments

We thank C. Kang (Seoul National University) for providing senolytic agents, Dasatinib and Quercetin.

## Data and materials availability

The bulk RNA sequencing data and scRNA-seq data have been deposited at Gene Expression Omnibus (GEO, GSE accession number: GSE254475 and GSE254476, respectively) and are publicly available as of the date of publication. Source data supporting the findings of this study are provided with this paper. Additional data that support the findings of this study are available from the corresponding author upon reasonable request.

