## Supplementary Figures and Legends for "Rejuvenation-Responsive and Senolytic-Sensitive Muscle Stem Cells Unveiled by CD200 and CD63 in Geriatric Muscle"

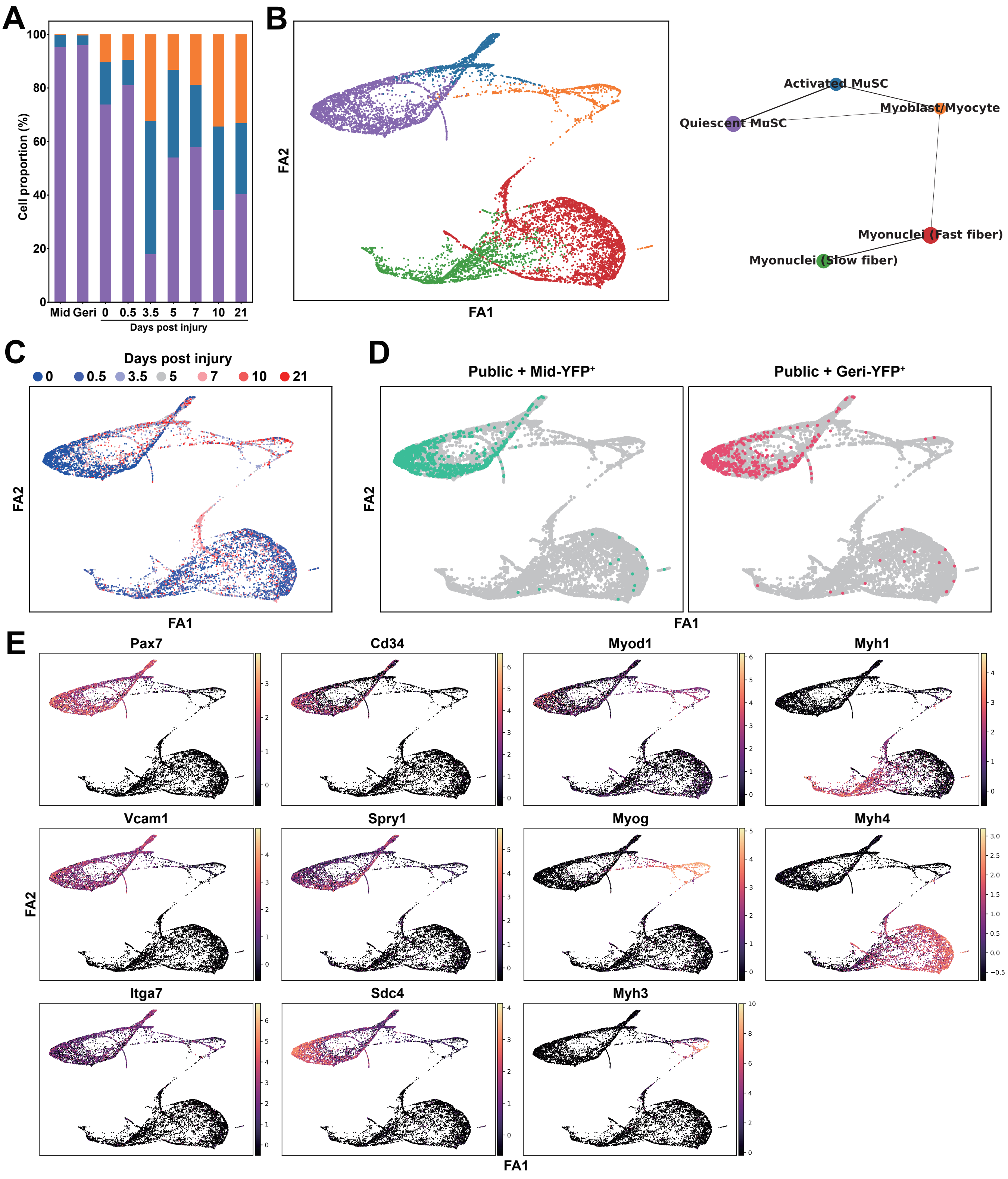

A
B
C
D
E
Cell proportion (%)
Mid Geri 0 0.5 3.5 5 7 10 21
Days post injury
FA1
FA2
Activated MuSC
Quiescent MuSC
Myoblast/Myocyte
Myonuclei (Fast fiber)
Myonuclei (Slow fiber)
Days post injury
0 0.5 3.5 5 7 10 21
Public + Mid-YFP+
Public + Geri-YFP+
Pax7
Cd34
Myod1
Myh1
Vcam1
Spry1
Myog
Myh4
Itga7
Sdc4
Myh3

**Fig. S1. Quiescent MuSC characteristics of YFP$^+$ cells.** (**A**) Stacked bar graph of myogenic cell cluster proportion determined in Fig. 2.at Mid, Geri, and each days-post-injury (DPI) of public data. (**B**) Force-directed graph (left) and partition-based graph abstraction (PAGA) connectivities (right) of YFP$^+$ cells integrated with public scRNA-seq data described in Fig. 2, colored by cell types. (**C**) Force-directed graph showing identities of cells based on the days-post-injury. (**D**) Force-directed graph showing the locations of YFP$^+$ cells of Mid and Geri in this study. (**E**) The expression levels of representative genes of specific cell types shown in the force-directed graph.

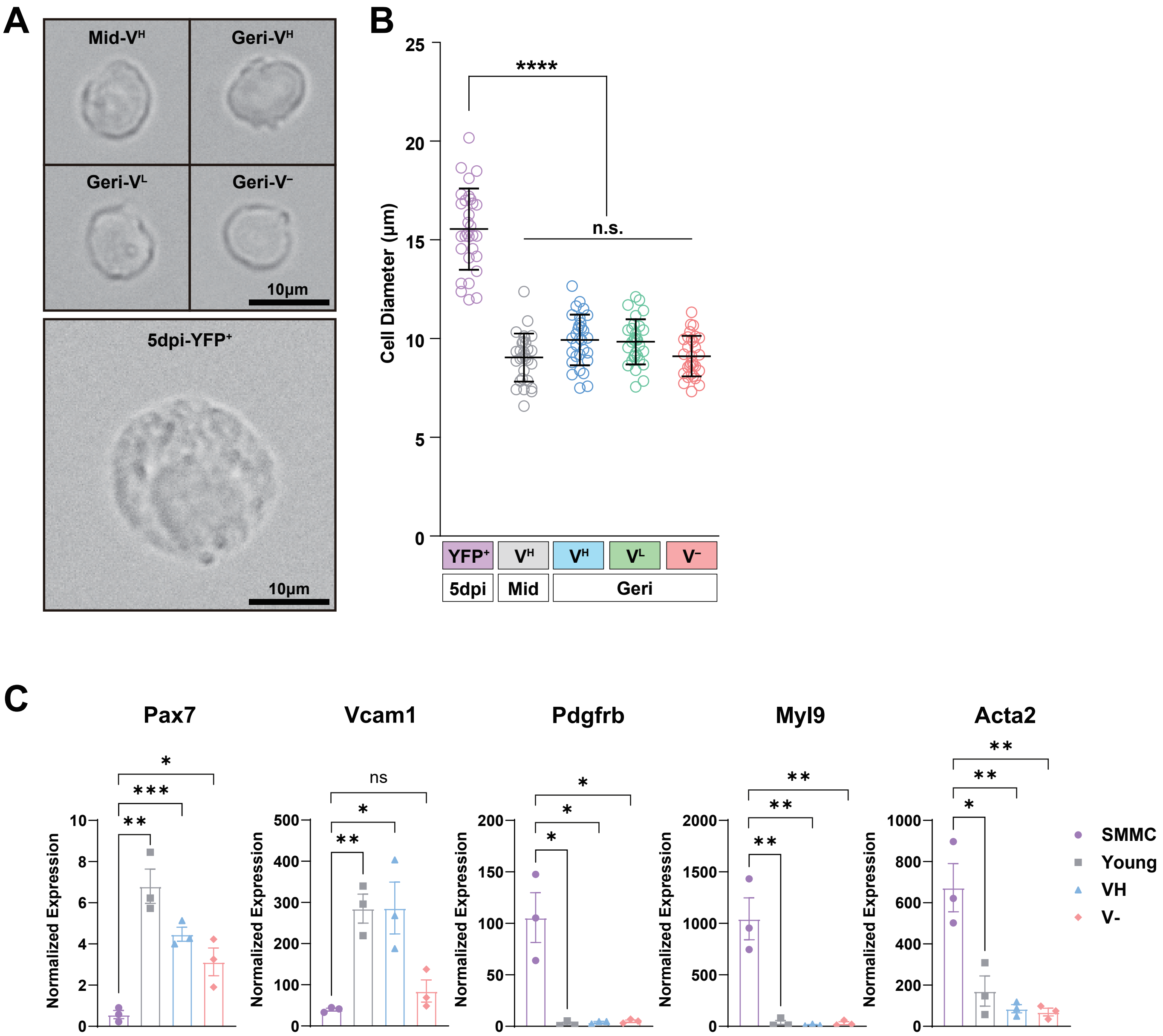

A
Mid-V^H
Geri-V^H
Geri-V^L
Geri-V^−
10μm
5dpi-YFP^+
10μm
B
****
n.s.
Cell Diameter (μm)
25
20
15
10
5
0
YFP^+
V^H
V^H
V^L
V^−
5dpi
Mid
Geri
C
Pax7
Vcam1
Pdgfrb
Myl9
Acta2
Normalized Expression
ns
SMMC
Young
VH
V-

**Fig. S2. VCAM1$^{-}$ cells resemble quiescent MuSCs in size and differ from SMMCs.** (**A**) Representative bright field image of freshly isolated YFP$^{+}$ cells of injured-5dpi young (20-weeks-old), Mid V$^{H}$, Geri V$^{H}$, V$^{L}$, and V$^{-}$. (**B**) Quantification of cell diameter in YFP$^{+}$ cells of injured-5dpi, Mid V$^{H}$, Geri V$^{H}$, V$^{L}$, and V$^{-}$, one-way ANOVA followed by Bonferroni's multiple comparison test; $p$ <0.0001 for injured-5dpi vs all YFP$^{+}$ cell groups of uninjured muscle. (**C**) Normalized expression of MuSC marker genes, Pax7 and Vcam1, and smooth muscle mesenchymal cells marker genes, Pdgfrb, Myl9 and Acta2.

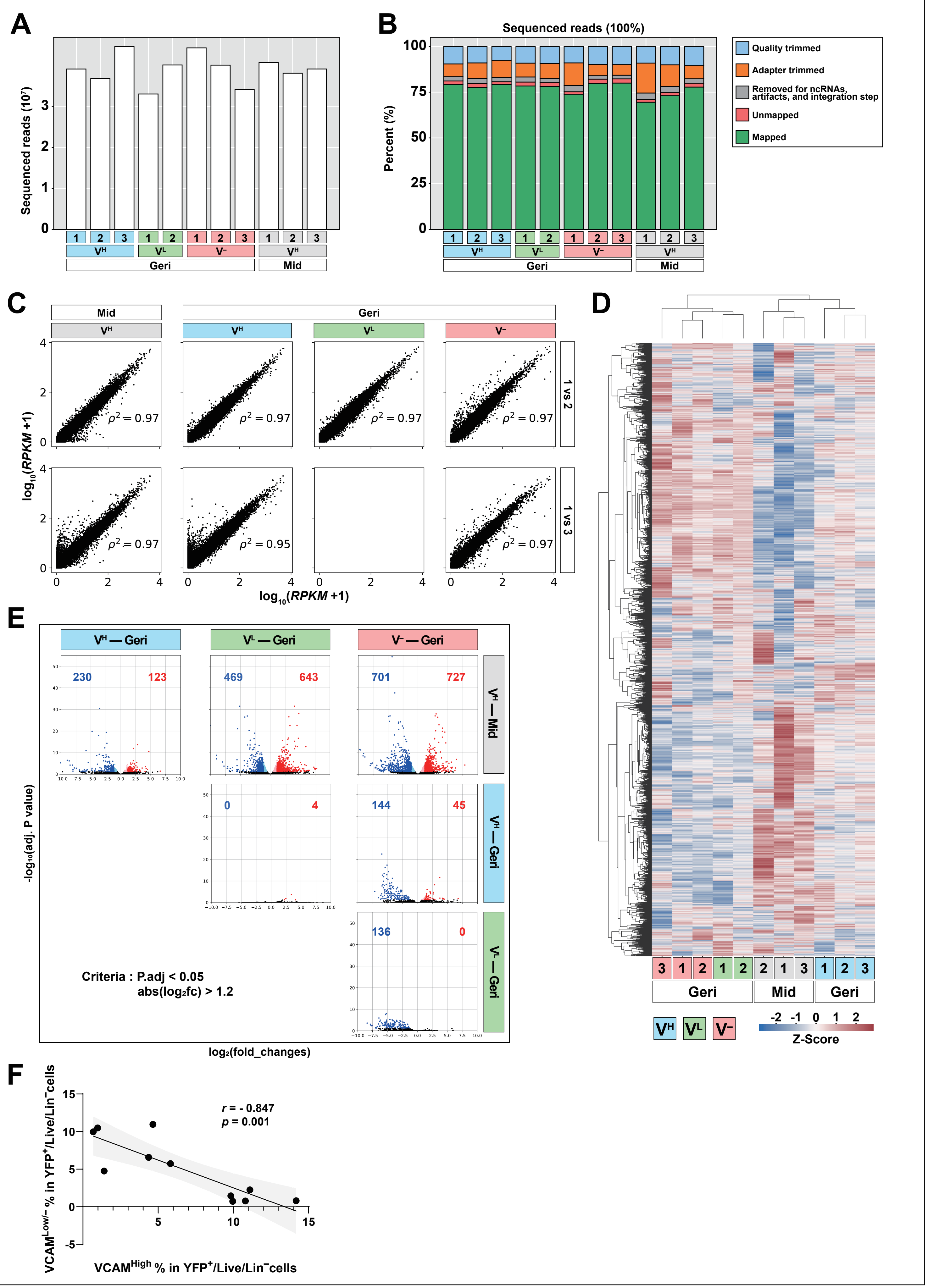
A
Sequenced reads (10^7)
VH VL V− VH
Geri Mid
B
Sequenced reads (100%)
Percent (%)
Quality trimmed
Adapter trimmed
Removed for ncRNAs, artifacts, and integration step
Unmapped
Mapped
C
Mid Geri
log10(RPKM +1)
1 vs 2
1 vs 3
D
Z-Score
E
VH — Geri
VL — Geri
V− — Geri
VH — Mid
VH — Geri
VL — Geri
230 123
469 643
701 727
0 4
144 45
136 0
-log10(adj. P value)
log2(fold_changes)
Criteria : P.adj < 0.05
abs(log2fc) > 1.2
F
r = - 0.847
p = 0.001
VCAMLow/− % in YFP+/Live/Lin−cells
VCAMHigh % in YFP+/Live/Lin−cells

**Fig. S3. Validation and basic analysis of transcriptomic data of YFP$^+$ cells.** (**A**) Total number of sequenced reads of each sample. (**B**) The percentage of trimmed reads of each sample for quality control. (**C**) Dot plots of gene expression showing linear correlations of between samples of the same group. The number in the plot represents Spearman Coefficient. (**D**) Hierarchical clustering of bulk RNA-seq samples with gene expression profiles, showing sample similarity (x-axis) and gene similarity (y-axis), illustrating inter-group similarity and similarity between $V^H$ of Mid and Geri and between $V^L$ and $V^–$ of Geri. (**E**) Volcano plots showing differentially expressed genes (DEGs) between groups at a threshold of LFC > 1.2 and adjusted *p* value < 0.05. The numbers in the plot represent the number of up-regulated (red) and down-regulated (blue) genes compared to the sample indicated in the right of the panel. (**F**) Correlation analysis between the percentages of $VCAM1^{High}$ and $VCAM1^{Low/–}$ cells within YFP$^+$/Live/Lin$^–$ MuSCs isolated from geriatric mice. Each dot represents an individual mouse. A significant negative correlation was observed (Pearson's $r = –0.847$, $p = 0.001$), indicating that a higher proportion of $VCAM1^{High}$ MuSCs is associated with a lower proportion of $VCAM1^{Low/–}$ MuSCs. Shaded area represents the 95% confidence interval of the linear regression.

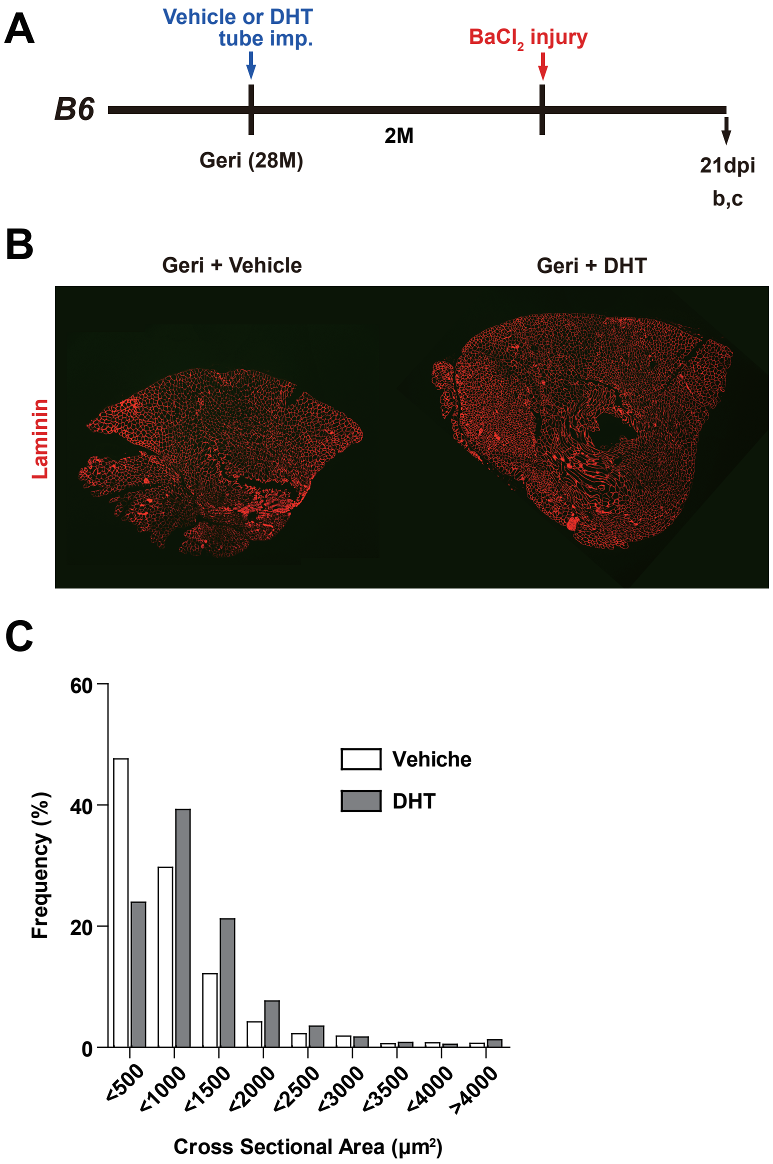
A
Vehicle or DHT
tube imp.
BaCl2 injury
B6
2M
Geri (28M)
21dpi
b,c
B
Geri + Vehicle
Geri + DHT
Laminin
C
60
40
20
0
Frequency (%)
Vehiche
DHT
<500
<1000
<1500
<2000
<2500
<3000
<3500
<4000
>4000
Cross Sectional Area (μm2)

**Fig. S4. Rehabilitation of myofiber regeneration capacity in geriatric mice after DHT supplementation.**

(**A**) Experimental schematics geriatric B6 mice with DHT or vehicle tube implantation before BaCl2 injury. (**B**) Representative images of immunohistochemistry staining for laminin in geriatric mice with vehicle or DHT tube implantation. (**C**) Quantification of the cross-sectional area of the 21 dpi TA muscle of geriatric mice after vehicle or DHT tube implantation.
